# Different combinations of ErbB receptor dimers generate opposing signals that regulate cell proliferation in cardiac valve development

**DOI:** 10.1101/067249

**Authors:** Ryo Iwamoto, Naoki Mine, Hiroto Mizushima, Eisuke Mekada

## Abstract

HB-EGF plays an indispensable role in suppression of cell proliferation in mouse valvulogenesis. However, ligands of the EGF receptor (EGFR/ErbB1), including HB-EGF, are generally considered as growth-promoting factors, as shown in cancers. HB-EGF binds to and activates ErbB1 and ErbB4. We investigated the role of ErbB receptors in valvulogenesis in vivo using ErbB1- and ErbB4-deficient mice, and an ex vivo model of endocardial cushion explants. We show that HB-EGF suppresses valve mesenchymal cell proliferation through a heterodimer of ErbB1 and ErbB4, and an ErbB1 ligand(s) promotes cell proliferation through a homodimer of ErbB1. Moreover, a rescue experiment with cleavable or uncleavable isoforms of ErbB4 in *ERBB4* null cells suggests that the cytoplasmic intracellular domain of ErbB4, rather than the membrane-anchored tyrosine kinase, achieves this suppression. Our study demonstrates that opposing signals generated by different ErbB dimer combinations function in the same cardiac cushion mesenchymal cells for proper cardiac valve formation.

**Summary statement:** In valvulogenesis, opposing signals generated by different combinations of ErbB-dimers elaborately regulate cell proliferation, in which proteolytically released intracellular domain of ErbB4 activated by HB-EGF is required to suppress proliferation.

## Introduction

The ErbB system, comprising epidermal growth factor (EGF) family members and ErbB receptor tyrosine kinases (RTKs), is fundamental in cell proliferation, differentiation, migration, and survival. In mammals, the EGF-ErbB system consists of four receptors: EGF receptor (EGFR)/ErbB1, ErbB2, ErbB3 and ErbB4. It also includes multiple EGF family ligands including EGF, transforming growth factor-alpha (TGF-α), amphiregulin (ARG), heparin-binding EGF-like growth factor (HB-EGF), betacellulin (BTC), epiregulin (ERG), epigen (EPG), and neuregulin (NRG) 1–4. Although EGF family ligands share a conserved receptor-binding motif, and soluble forms are usually produced from membrane-anchored precursors (Harris et al., 2003), EGF family members bind to the various ErbB receptors with differing degrees of preference (Riese and Stern, 1998). Moreover, in addition to the different ligand specificities and downstream signalling molecules of ErbB receptors, they form homo- or hetero-dimers in various combinations (Olayioye et al., 2000). Thus, the vast number of ligand and receptor combinations provides redundancy and robustness in the EGF-ErbB system, which makes it possible for the bioactivities exerted by the system to be finely tuned. The complexity of the EGF-ErbB system implies that different combinations of EGF family ligands and ErbB RTKs achieve a range of cellular behaviours in vivo. However, the physiological significance of such diversity in the intracellular signals generated by the EGF-ErbB system remains to be elucidated. Moreover, there is little information about how such diverse signals are precisely regulated in vivo, and which EGF ligand-ErbB RTK and ErbB dimer combinations specifically function in certain physiological processes. In this study, we aimed to resolve these issues.

HB-EGF binds to ErbB1 and ErbB4 (Higashiyama et al., 1991; Elenius et al., 1997; Raab and Klagsbrun, 1997). Analyses of HB-EGF null mice have revealed that HB-EGF plays pivotal roles in several physiological and pathological processes (Mekada and Iwamoto, 2008). Similar to other ErbB1 ligands, because HB-EGF is a potent mitogen that has been implicated in cancer progression and malignancy, it is generally known as a growth-promoting factor. However, in developmental and physiological processes, HB-EGF is not characterised as a growth-promoting factor but rather acts as a growth-inhibiting or migration-promoting factor (Mekada and Iwamoto, 2008). Cardiac valve development (valvulogenesis) is an example of a process in which HB-EGF functions as a growth-inhibiting factor (Iwamoto and Mekada, 2006).

Mouse valvulogenesis occurs in two consecutive steps: cardiac cushion formation and valve remodelling (Schroeder et al., 2003; Armstrong and Bischoff, 2004). The cardiac cushion is the primordial tissue of cardiac valves, and the cushion formation stage is accompanied by endothelial-to-mesenchymal transition. In mice, this stage is completed by embryonic day (E) 12.5 (Lakkis and Epstein, 1998) and is followed by remodelling of the cushions to form thin valve leaflets. Although much is known about the signalling pathways involved in cushion formation, the molecular mechanisms underlying valve remodelling are still poorly understood. HB-EGF is expressed and functions during the remodelling process (Iwamoto et al., 2010). Several HB-EGF-deficient mouse lines show valve abnormalities, indicating that HB-EGF is critical for normal valvulogenesis (Iwamoto et al., 2003; Jackson et al., 2003; Yamazaki et al., 2003; Iwamoto and Mekada, 2006; Iwamoto et al., 2010). Such mutant mice have enlarged cardiac valves due to hyperproliferation of mesenchymal cells. These studies indicate that the soluble form of HB-EGF, which is secreted from endocardial cells, functions as a ‘growth-inhibiting’ factor for mesenchymal cells. In addition, they suggest that signals promoting cell proliferation exist in parallel with the HB-EGF-mediated inhibitory cascade. However, the precise molecular mechanism controlling such mesenchymal cell proliferation is still unclear.

To address these issues, we performed in vivo experiments using several mutant mouse lines and ex vivo experiments using cardiac cushion explants. The ex vivo experimental culture system recapitulates in vivo phenomena and has shown a functional relationship between HB-EGF and mesenchymal cell proliferation (Iwamoto et al., 2010). In the present study, we demonstrate the regulatory mechanism for mouse valvulogenesis, in which ErbB RTKs finely regulate mesenchymal cell proliferation in a dual mode via dimerization with different partners: ErbB1 and ErbB4 form a heterodimer receptor for HB-EGF, resulting in growth inhibition, whereas homodimerization of ErbB1 produces a receptor for a certain ErbB1 ligand(s), leading to growth promotion. Moreover, in the former inhibitory signalling, proteolytic processing of ErbB4 is required to release the intracellular domain (ICD) of ErbB4. These results provide evidence of a novel and elaborate mechanism for the regulation of cell proliferation by specific ErbB dimer combinations in vivo.

## Results

### ErbB1 is involved in HB-EGF-mediated suppression of valve mesenchymal cell proliferation, as well as ErbB1 ligand-mediated up-regulation of proliferation in the absence of HB-EGF

To investigate the molecular mechanism governing HB-EGF-associated regulation of cell proliferation in valvulogenesis, we performed ex vivo experiments using cardiac cushion explants (Iwamoto et al., 2010). In this culture system, a cushion explant from an E10.5 embryonic heart was placed on a type I collagen gel containing hyaluronic acid (HA-Col) (Fig. 1A). Endocardial cells derived from the endocardium of the cushion explants (ON-gel cells) proliferated and spread on the surface of the HA-Col, while some endocardial cells differentiated to mesenchymal cells that invaded into the HA-Col and proliferated (IN-gel cells). Under this culture condition, IN-gel cells from HB-EGF knockout (KO) (*HB^del/del^*) explants proliferated much more than those from wild-type (WT) explants, whereas ON-gel cells from WT and *HB^del/del^* cushions proliferated at comparable levels (Fig. 1B,C). Therefore, this ex vivo experimental system reproduced in vivo data and showed a functional relationship between HB-EGF and mesenchymal cell proliferation (Iwamoto et al., 2010). Up-regulation of the mesenchymal cell proliferation of *HB^del/del^* explants suggests that HB-EGF functions as a ‘growth-inhibiting’ factor for mesenchymal cells, and that an unknown signal(s) promoting cell proliferation exists in parallel with the HB-EGF-mediated inhibitory cascade. Thus, we investigated which receptors transmit each signal in parallel by first focusing on ErbB1. ErbB1 is thought to be involved in the inhibitory signals induced by HB-EGF, because *ERBB1* null mice develop cardiac valve enlargement as observed in *HBEGF* null mice (Chen et al., 2000; Jackson et al., 2003).

**Fig. 1.**
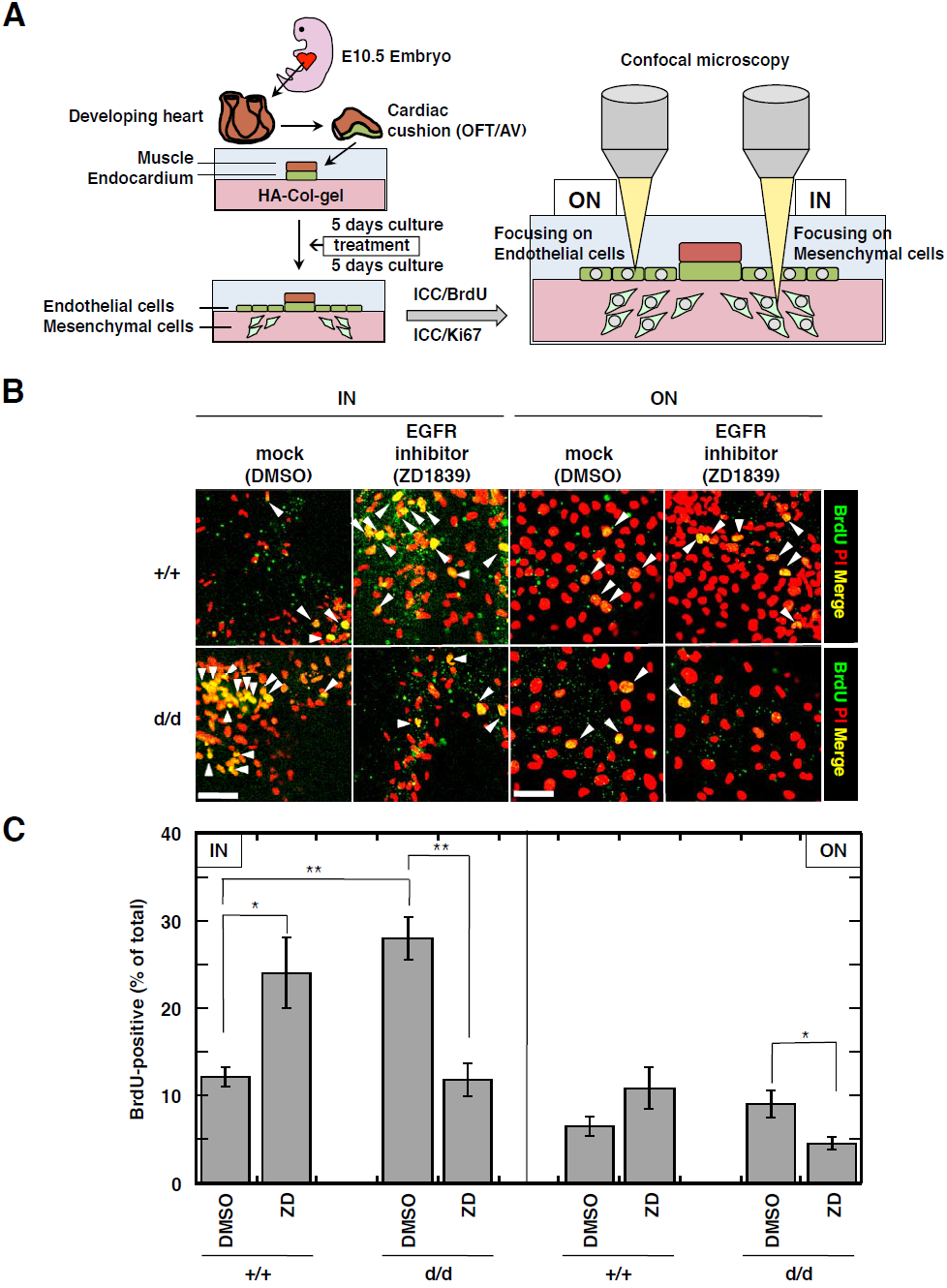
Inhibition of ErbB1 in cushion explants. **(A)** Schematic illustration of the cushion explant culture system. **(B)** Representative images of mesenchymal (IN) and endocardial (ON) cells of cushion explants from WT (+/+) and *HB^del/del^* (d/d) embryos that were mock treated (DMSO) or treated with ZD1839 (0.3 μM), an EGFR inhibitor. Cells were stained with propidium iodide (PI) (red, nuclei) and an anti-BrdU antibody (green), and then the images were merged. Yellow cells represent BrdU-positive cells (arrowheads). Bar, 50 μm for all panels. **(C)** Scoring of BrdU-positive mesenchymal (IN) and endocardial (ON) cells of WT (+/+) and *HB^del/del^* (d/d) cushion explants treated with ZD1839 based on the data shown in B. ZD, ZD1839. Data represent the mean values ± SE of results obtained from at least five individual explants; n = 6, 5, 9, and 5 for DMSO:+/+, ZD:+/+, DMSO:d/d, and ZD:d/d, respectively. \**p* < 0.05; \*\**p* < 0.01. See also Fig. S1.

First, we examined ZD1839, a chemical inhibitor of ErbB1 (Fig. 1B,C). ZD1839 up-regulated WT cell proliferation as observed previously (Iwamoto et al., 2010). Moreover, surprisingly, ZD1839 down-regulated hyper-proliferation of *HB^del/del^* cells. These results strongly suggest that ErbB1 activity is required not only for HB-EGF-mediated suppression of WT cell proliferation, but also for promotion of increased proliferation in *HB^del/del^* cells.

To confirm the above possibility in vivo, we used a hypomorphic ErbB1 mutant, *waved-2* (Luetteke et al., 1994), and tested for a genetic interaction between *HBEGF* and *ERBB1*. In *waved-2* mice, the kinase activity of ErbB1 is decreased to <10% of that of WT ErbB1 because of a point mutation in the kinase domain. Although *waved-2* single mutant embryos (+/+; *wa2*/*wa2*) showed a weak but significant valve-enlarged phenotype, the abnormalities in *HB^del/del^* and *waved-2* double mutant embryos (*d/d; wa2/wa2*) were significantly weaker than those of *HB^del/del^* single mutants (*d/d; +/+*) (Fig. 2). These results indicate that ErbB1 is also required to promote the increased proliferation of *HB^del/del^* cells. Immunohistochemistry for phospho-ErbB1 showed that the phosphorylation rate of ErbB1 was comparable between WT and *HB^del/del^* valves (Fig. S1). This finding suggests that, even in the condition without HB-EGF, a certain ligand(s) of ErbB1 still activates ErbB1 in mesenchymal cells of developing valves. Taken together, these results indicate that ErbB1 functions in both HB-EGF-mediated inhibition of mesenchymal cell proliferation and promotion of hyperproliferation in HB-EGF-lacking cells.

**Fig. 2.**
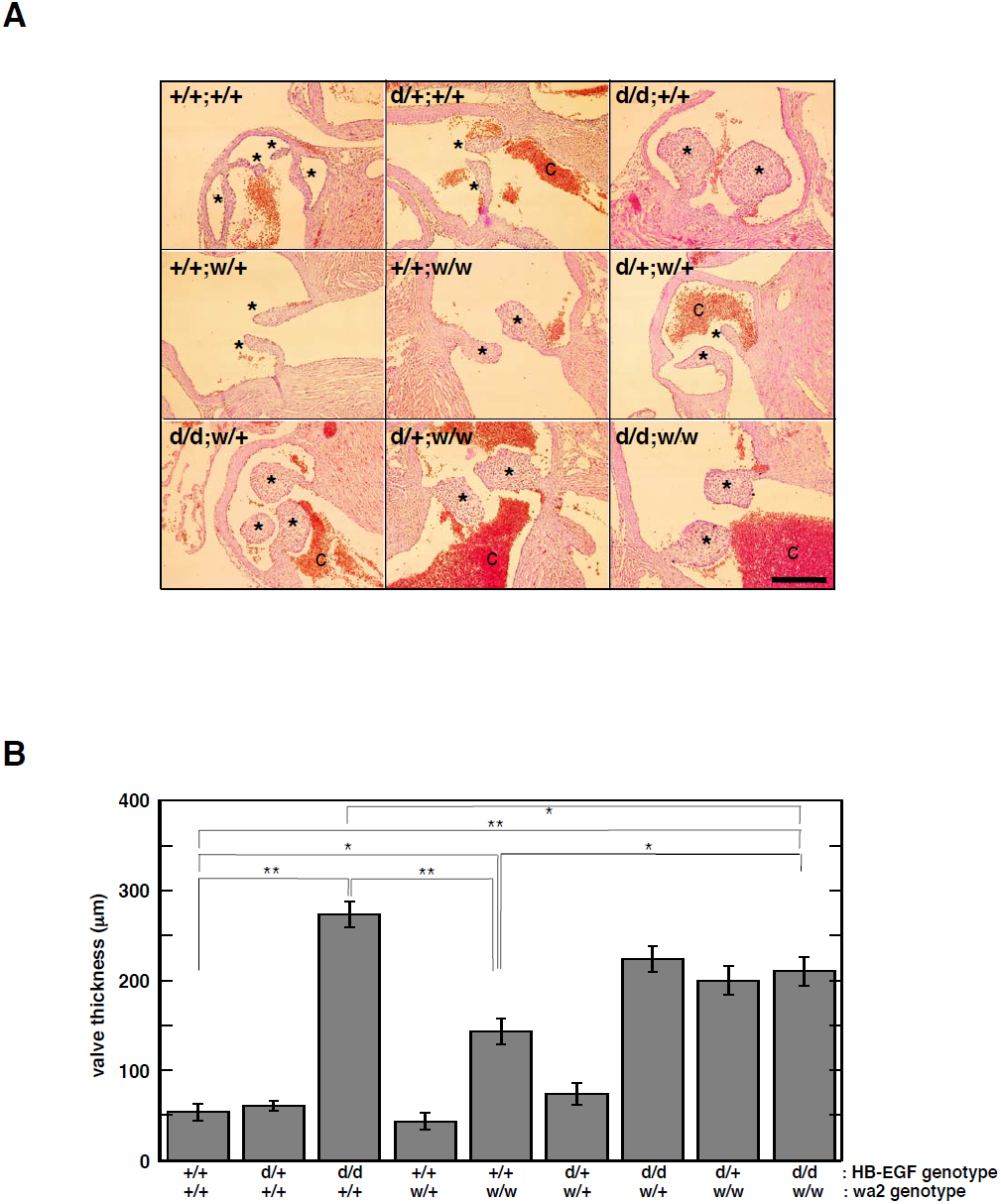
Decreased cardiac valve enlargements in a double mutant embryo carrying *HB^de^* and *waved-2* EGFR alleles. **(A)** Representative images of cardiac valves. Haematoxylin and eosin-stained longitudinal sections of E19.5 embryo hearts from mice of the indicated genotype (+/+, WT; d/+, *HB^del/+^; d/d, HB^del/del^*; *w/+, waved-2/+; w/w; waved-2/waved-2*) are shown. Pulmonic or aortic valves are indicated by asterisks. Blood clots are indicated as ‘C’. Bar, 200 μm for all panels. **(B)** Comparison of pulmonic or aortic valve thicknesses among E19.5 embryo hearts from mice of the indicated genotype based on the data shown in A. The largest diameter of each valve in serial sections was measured. The valve sizes were calculated as the mean ± SE of results obtained from three individual embryos for all points. \**p* < 0.05; \*\**p* < 0.01. See also Fig. S1.

### ErbB4 is involved in HB-EGF-mediated suppression of valve mesenchymal cell proliferation

Regarding the mechanism of ErbB1 signalling in both inhibition and promotion of cell proliferation, we hypothesised that ErbB1 may function by forming a dimer with different ErbB partners to achieve each function. To examine this possibility, we checked the expression of each ErbB receptor mRNA in explanted cells. RT-PCR analysis showed expression of ErbB1 and ErbB4 but little or no expression of ErbB2 and ErbB3 in valve cells (Fig. S2A). This result suggests that ErbB1 may function by forming a dimer with ErbB1 itself (ErbB1/ErbB1 homodimer) and/or with ErbB4 (ErbB1/ErbB4 heterodimer).

Next, we examined the effect of forced expression of a dominant-negative form of ErbB1 (dn-ErbB1) (Kashles et al., 1991) (Fig. 3A, Fig. S2B). Similar to the effect of the inhibitor, forced expression of dn-ErbB1 not only up-regulated WT mesenchymal cell proliferation but also down-regulated hyperproliferation of *HB^del/del^* cells (Fig. 3C,D). These results confirmed that ErbB1 functions in both HB-EGF-mediated inhibition of mesenchymal cell proliferation and promotion of hyperproliferation in the absence of HB-EGF. Next, we examined dn-ErbB2 (Jones and Stern, 1999), dn-ErbB3 (Ram et al., 2000), and dn-ErbB4 (Jones et al., 1999) (Fig. 3A, Fig. S2B) to determine which ErbB receptor functions with ErbB1 in each process. Consistent with their low endogenous expression, dn-ErbB2 and dn-ErbB3 had little effect on mesenchymal cell proliferation. Conversely, dn-ErbB4 increased cell proliferation by inhibiting HB-EGF-mediated suppression of WT mesenchymal cell proliferation, but not hyperproliferation of *HB^del/del^* cells (Fig. 3C,D). These results strongly suggest the specific involvement of ErbB4 in the HB-EGF-mediated inhibitory process. The dn-ErbB4 used here was a deletion mutant lacking the cytoplasmic domain (Fig. 3A). Thus, there was the possibility that dn-ErbB4 depletes HB-EGF in the culture system, and the effect of the dominate-negative mutant did not represent the supposed dominant-negative effect of ErbB4. However, addition of excess amounts of recombinant HB-EGF protein did not alter the effects of dn-ErbB4 (Fig. S3), ruling out the possibility of HB-EGF depletion by dn-ErbB4. Taken together, these results strongly suggest that activation of the heterodimer consisting of ErbB1 and ErbB4 by HB-EGF suppresses cell proliferation, whereas activation of the ErbB1 homodimer by ErbB1 ligands promotes cell proliferation.

**Fig. 3.**
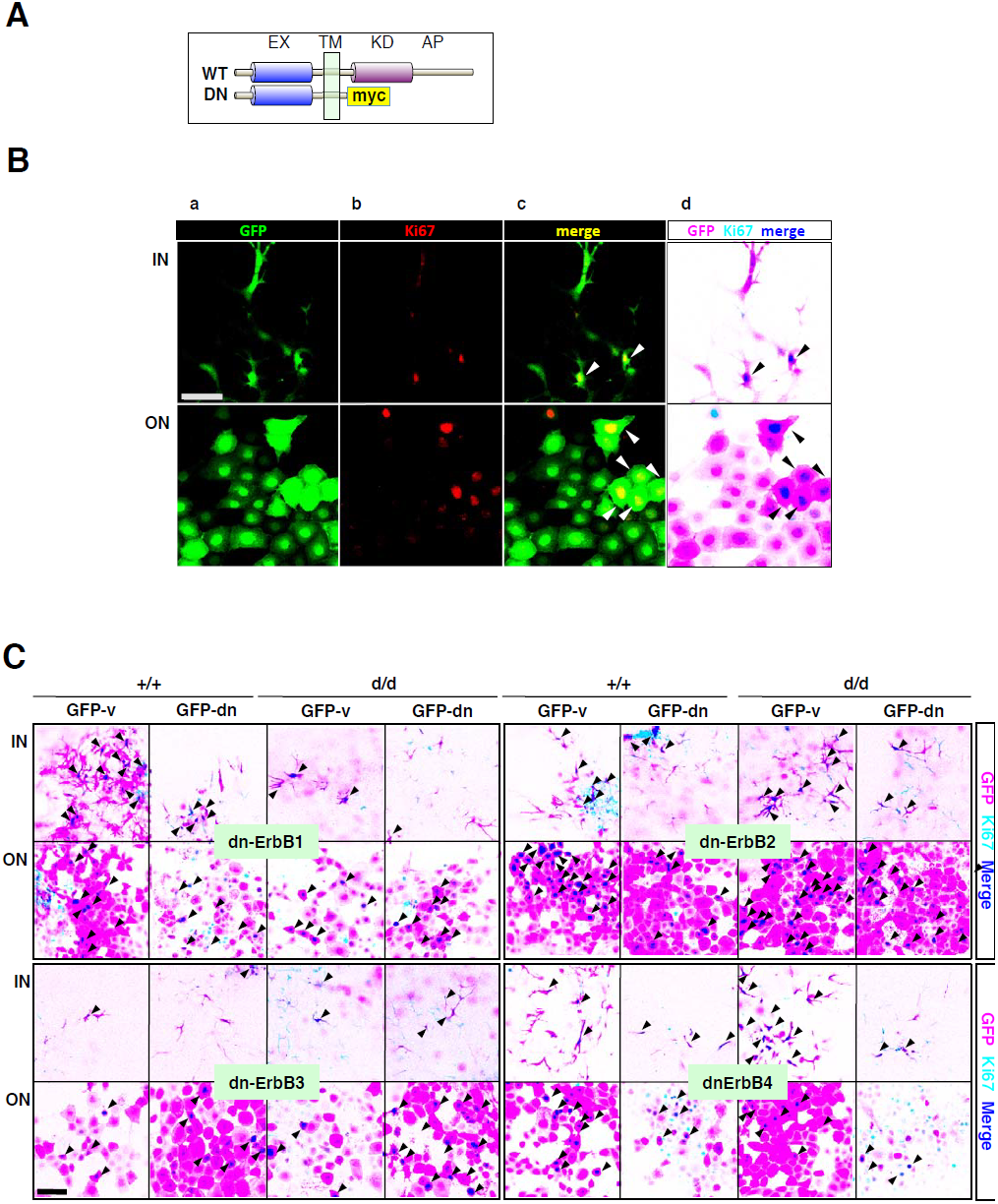
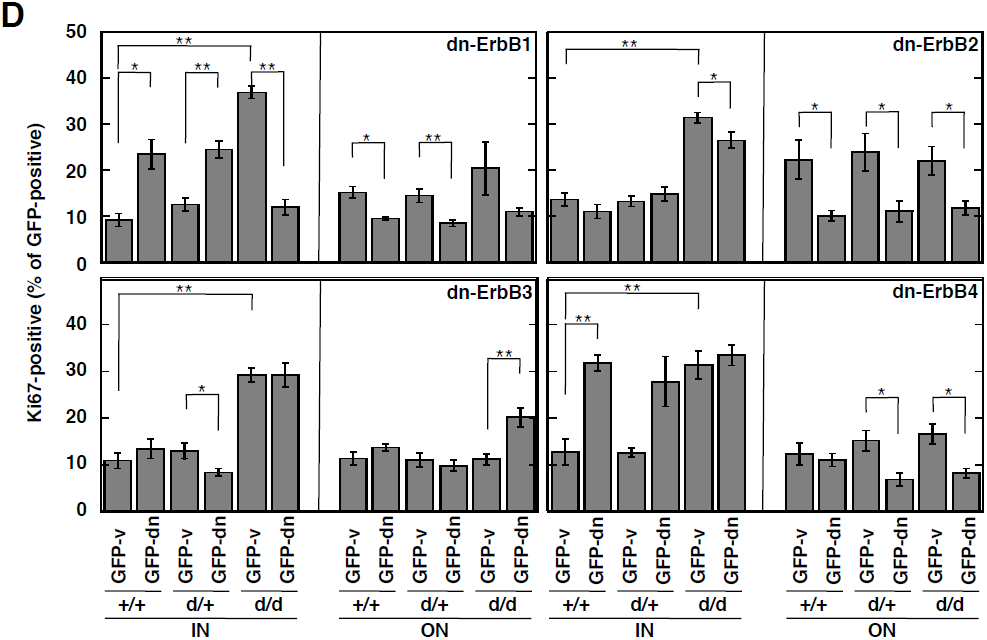
Effects of the expression of a dominant-negative mutant for each ErbB receptor on cell proliferation in cushion explants. **(A)** Schematic representation of wild-type ErbB1–4 (WT) and dominant-negative forms of ErbB1–4 with a myc tag (DN). EX, extracellular domain; TM, transmembrane domain; KD, kinase domain; AP, autophosphorylation domain; myc, myc tag. **(B)** Representative images of mesenchymal (IN) and endocardial (ON) cells of cushion explants from WT embryos that were transfected with an empty GFP vector as shown in green (a). Cells were stained with an anti-Ki67 antibody as shown by red nuclei (b), and then the images were merged with GFP images (c). Green cells with yellow nuclei represent Ki67-positive cells (arrowheads). However, green fluorescence of the whole cell was too strong to distinguish the yellow nucleus. Thus, to obtain clearer double-positive cells with yellow nuclei from GFP single-positive cells with a green whole cell body, the image (c) was automatically colour converted [d, each colour converted as green to magenta (GFP), red to pale blue (Ki67), and yellow to indigo blue (merge)] by image software as described in the “Experimental Procedures”. Thus in (d), magenta cells with an indigo blue nucleus represent GFP and Ki67 double-positive cells (arrowheads). Bar, 50 μm for all panels. **(C)** Representative images of mesenchymal (IN) and endocardial (ON) cells of cushion explants from WT (+/+) and *HB^del/del^* (d/d) embryos that were mock transfected with an empty GFP vector (GFP-v) or each dominant-negative mutant IRES-linked GFP vector (GFP-dn) for ErbB1 (dn-ErbB1), ErbB2 (dn-ErbB2), ErbB3 (dn-ErbB3), and ErbB4 (dn-ErbB4). Cells were stained with an anti-Ki67 antibody, and then the images were merged with GFP images. All images were automatically colour converted. Magenta cells with an indigo blue nucleus represent GFP and Ki67 double-positive cells (arrowheads). Bar, 100 μm for all panels. **(D)** Scoring of GFP and Ki67 double-positive mesenchymal (IN) and endocardial (ON) cells of WT (+/+), *HB^del/+^* (d/+), and *HB^del/del^* (d/d) cushion explants transfected with each dominant-negative mutant based on the data shown in C. Data represent the mean values ± SE of results obtained from at least three individual explants; dn-ErbB1, n = 3, 4, 6, 6, 3, and 3 for GFP-v-+/+, GFP-dn-+/+, GFP-v-d/+, GFP-dn-d/+, GFP-v-d/d, and GFP-dn-d/d, respectively; dn-ErbB2, n = 6, 3, 5, 7, 6, and 5 for GFP-v:+/+, GFP-dn:+/+, GFP-v:d/+, GFP-dn:d/+, GFP-v:d/d, and GFP-dn:d/d, respectively; dn-ErbB3, n = 6, 5, 6, 3, 5, and 6 for GFP-v:+/+, GFP-dn:+/+, GFP-v:d/+, GFP-dn:d/+, GFP-v:d/d, and GFP-dn:d/d, respectively; dn-ErbB4, n = 4, 4, 6, 3, 5, and 5 for GFP-v:+/+, GFP-dn:+/+, GFP-v:d/+, GFP-dn:d/+, GFP-v:d/d, and GFP-dn:d/d, respectively. *p < 0.05; \*\**p* < 0.01. See also Fig. S2 and Fig. S3.

### Up-regulation of valve mesenchymal cell proliferation in ErbB4 KO mice

To investigate whether ErbB4 regulates growth inhibition during valvulogenesis in vivo, we analysed the valve phenotype of *ERBB4-KO (ERBB4^−/−^*) mice and examined the proliferation of mesenchymal cells in ex vivo cultures. Because of the early embryonic lethality in *ERBB4^−/−^* mice before cushion formation (Gassmann et al., 1995), which is due to a defect in heart muscle trabeculation, we analysed a mutant mouse line rescued from early embryonic lethality by reintroduction of human ErbB4 (HER4) in cardiomyocytes (Tidcombe et al., 2003). Because the HER4 cDNA is under the control of the cardiac-specific α-myosin heavy chain (MHC) promoter, these mice express HER4 specifically in cardiomyocytes and not in any other cell type. Thus, the heart develops normally and these mice are born, reach adulthood, and are fertile (Tidcombe et al., 2003). Although the degree of abnormality was relatively milder, enlargement of the valves in heart-rescued *ERBB4^−/−^ (hr-ERBB4^−/−^*) newborns was phenotypically similar to that in *HB^del/del^* mice (Fig. S4). The mesenchymal cells of explants from *hr-ERBB4^−/−^* valves showed an approximately 2-fold higher proliferation rate compared with those from WT and *hr-ERBB4^+/−^* mice with statistical significance (Fig. 4). These results indicate that ErbB4 functions in HB-EGF-induced inhibition of mesenchymal cell proliferation in developing valves.

**Fig. 4.**
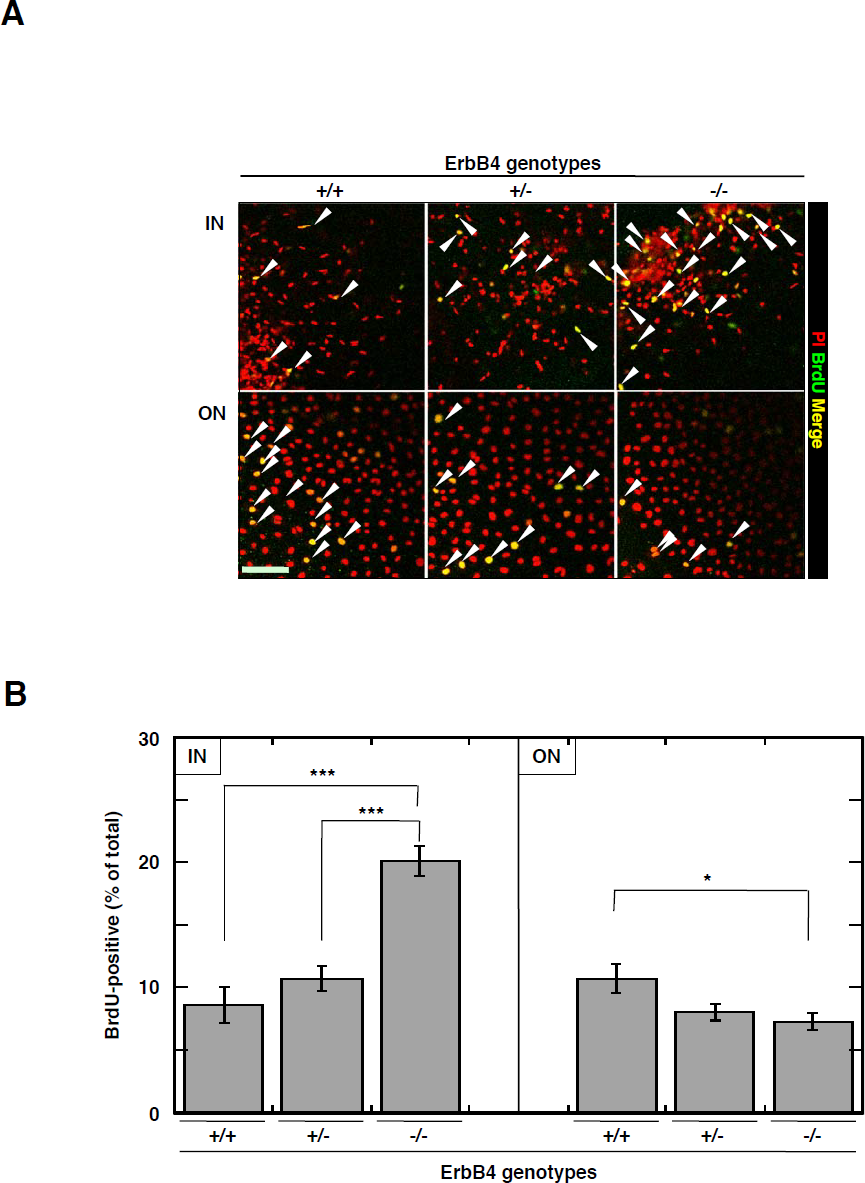
Increased proliferation of mesenchymal cells from *ERBB4* KO cushion explants. **(A)** Representative images of mesenchymal (IN) and endocardial (ON) cells of explants from WT (+/+), *ERBB4^+/−^* (+/−), and *ERBB4^−/−^* (−/−) embryos. Cells were stained with PI (red, nuclei) and an anti-BrdU antibody (green), and the images were merged. Yellow cells represent BrdU-positive cells (arrowheads). Bar, 100 μm for all panels. **(B)** Scoring of BrdU-positive mesenchymal (IN) and endocardial (ON) cells of explants from WT (+/+), *ERBB4^+/−^* (+/−), and *ERBB4^−/−^* (−/−) embryos based on the data shown in A. Data represent the mean values ± SE of results obtained from at least five individual explants; n = 6, 5, and 10 for +/+, +/−, and −/−, respectively. \**p* < 0.05; \*\*\**p* < 0.001. See also Fig. S4.

### A cleavable JM-a isoform of ErbB4 functions in the suppression of valve mesenchymal cell proliferation

Unlike other ErbB family RTKs, ErbB4 consists of alternatively spliced variants. Among them, ErbB4 is classified as JM-a or JM-b, if it is with or without a cleavage site by ADAM (a disintegrin and metalloproteinase) 17 in the juxtamembrane domain, respectively (Carpenter, 2003; Veikkolainen et al., 2011). When activated by ligand binding, the ectodomain of the JM-a type is proteolytically cleaved by ADAM17, generating the membrane-associated C-terminal fragment of ErbB4 (ErbB4-CTF). This cleavage is followed by intramembrane proteolysis by presenilin-1 (PS1), resulting in the translocation of the soluble intracellular domain of ErbB4 (ErbB4-ICD) into the nucleus (Lee et al., 2002; Medina and Dotti, 2002). The ICD then activates or inhibits the expression of several genes (Veikkolainen et al., 2011). Conversely, the JM-b type is uncleavable because of a difference in the amino acid sequence corresponding to the juxtamembrane domain, which is recognised by ADAM17. Thus, similar to other ErbB RTKs, the JM-b type activates canonical signalling cascades (Veikkolainen et al., 2011).

To reveal which ErbB4 isoform (JM-a or JM-b) dimerizes with ErbB1 to inhibit cell proliferation, we examined the expression of these isoforms at the mRNA level. RT-PCR analysis using specific primer sets for the mRNA for each isoform of *ERBB4* showed expression of both JM-a and JM-b in explanted cells (Fig. S5A). Next, we performed a rescue experiment by exogenous introduction of a vector encoding cleavable JM-a, uncleavable JM-b, or the cleaved soluble ICD (Fig. 5A–C) into explants of *ERBB4^−/−^* and *ERBB4^+/−^* values. As a result, cleavable JM-a significantly suppressed hyperproliferation of *ERBB4^−/−^* mesenchymal cells. Thus, cleavable JM-a rescued *ERBB4^−/−^* cells from the abnormal proliferation. Uncleavable JM-b did not affect the hyperproliferation of *ERBB4^−/−^* mesenchymal cells, but significantly increased the proliferation of *ERBB4^+/−^* cells by exerting a dominant-negative effect on HB-EGF-mediated suppression of cell proliferation. Expression of only the soluble ICD weakly but significantly suppressed hyperproliferation of *ERBB4^−/−^* mesenchymal cells (Fig. 5B,C). Similar results were obtained using explants of *HB^del/del^* and WT valves (Fig. 6). Cleavable JM-a significantly suppressed hyperproliferation of *HB^del/del^* mesenchymal cells, indicating that JM-a type ErbB4 functions downstream of HB-EGF. Uncleavable JM-b significantly increased the proliferation of WT cells, but did not affect hyperproliferation of *HB^del/del^* cells (Fig. 6). Taken together, these results indicate that JM-a type ErbB4 functions in this process, and suggest that the release of the ErbB4 ICD is required for the HB-EGF-mediated suppression of cell proliferation.

**Fig. 5.**
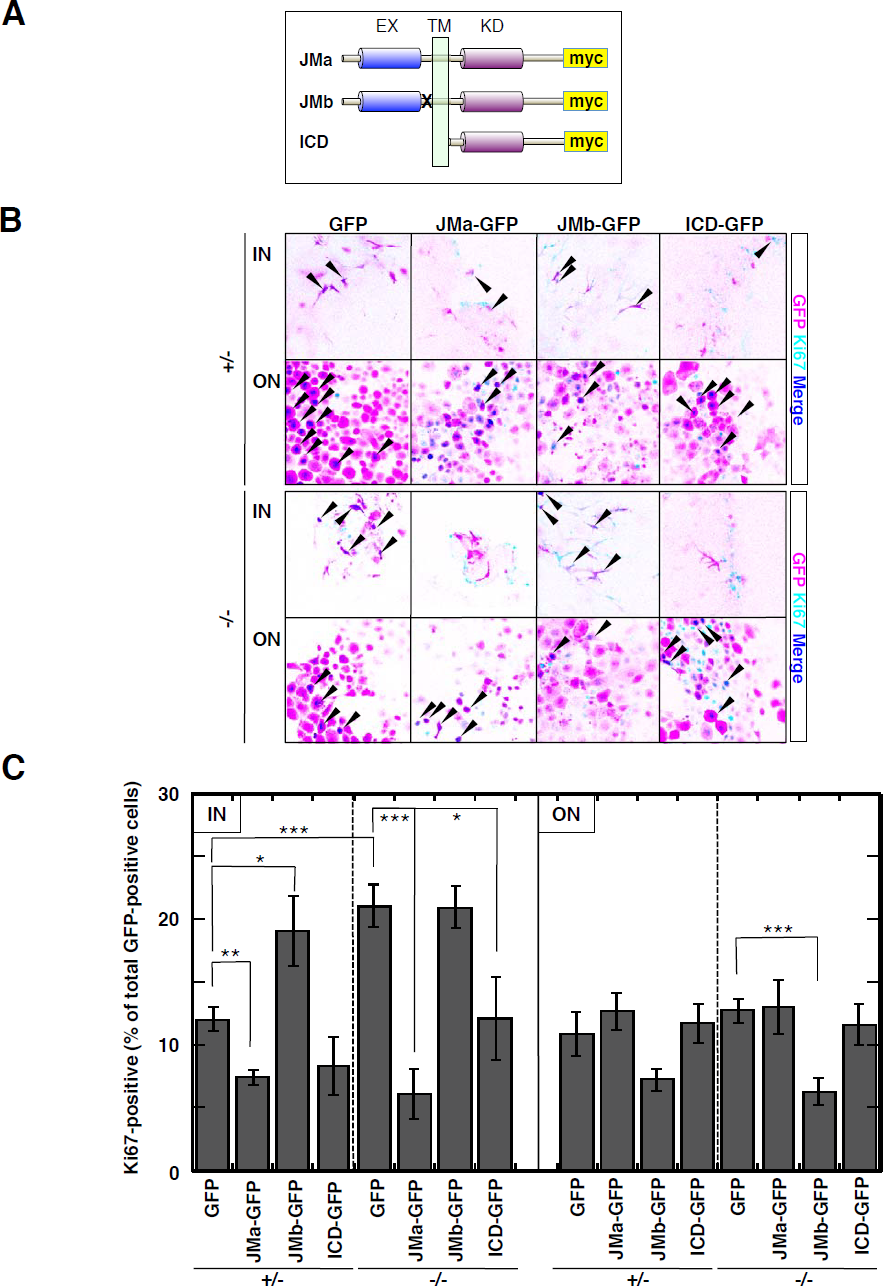
Rescue of *ERBB4* KO cells by each ErbB4 JM isoform or the ICD from abnormally increased proliferation in cushion explants. **(A)** Schematic representation of JM-a type ErbB4 with a myc tag (JMa), JM-b type ErbB4 with a myc tag (JMb), and only the ICD with a myc tag (ICD). EX, extracellular domain; TM, transmembrane domain; KD, kinase domain; X, uncleavable juxtamembrane site. **(B)** Representative images of mesenchymal (IN) and endocardial (ON) cells of cushion explants from *ERBB4^+/−^* (+/−) or *ERBB4^−/−^* (−/−) embryos that were mock transfected with an empty GFP vector (GFP), JM-a type ErbB4 with a myc tag IRES-linked GFP vector (JMa-GFP), JM-b type ErbB4 with a myc tag IRES-linked GFP vector (JMb-GFP), or the ICD of ErbB4 with a myc tag IRES-linked GFP vector (ICD-GFP). Cells were stained with an anti-Ki67 antibody, and then the images were merged with GFP images. All images were automatically colour converted. Magenta cells with an indigo blue nucleus represent GFP and Ki67 double-positive cells (arrowheads). Bar, 100 μm for all panels. **(C)** Scoring of GFP and Ki67 double-positive mesenchymal (IN) and endocardial (ON) cells of cushion explants from *ERBB4+* (+/−) and *ERBB4^−/−^* (−/−) embryos transfected with each construct based on the data shown in B. Data represent the mean values ± SE of results obtained from at least four individual explants; n = 7, 4, 5, 4, 9, 5, 8, and 4 for GFP:+/−, JMa-GFP:+/−, JMb-GFP:+/−, ICD-GFP:+/−, GFP:−/−, JMa-GFP:−/−, JMb-GFP:−/−, and ICD-GFP:−/−, respectively. \**p* < 0.05; \*\**p* < 0.01; \*\*\**P* <0.001. See also Fig. S5.

**Fig. 6.**
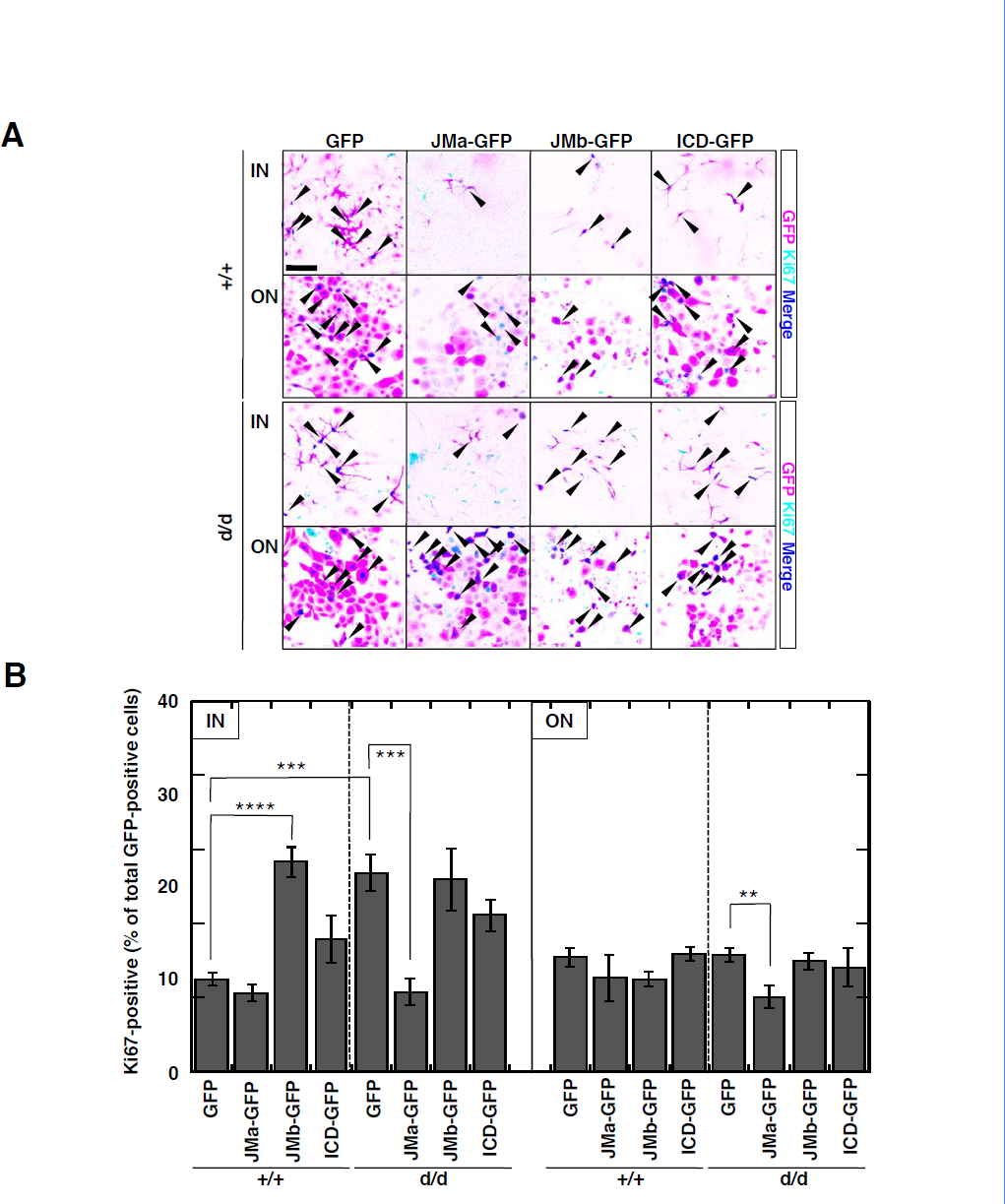
Effects of forced expression of each ErbB4 isoform or the ICD on abnormally increased proliferation of cells in *HB^del/del^* cushion explants. **(A)** Representative images of mesenchymal (IN) and endocardial (ON) cells of cushion explants from WT (+/+) and *HB^del/del^* (d/d) embryos that were mock transfected with an empty GFP vector (GFP), JM-a type ErbB4 with a myc tag IRES-linked GFP vector (JMa-GFP), JM-b type ErbB4 with a myc tag IRES-linked GFP vector (JMb-GFP), or the ICD of ErbB4 with a myc tag IRES-linked GFP vector (ICD-GFP). Cells were stained with an anti-Ki67 antibody, and then the images were merged with GFP images. All images were automatically colour converted. Magenta cells with an indigo blue nucleus represent GFP and Ki67 double-positive cells (arrowheads). Bar, 100 μm for all panels. **(B)** Scoring of GFP and Ki67 double-positive mesenchymal (IN) and endocardial (ON) cells of cushion explants from WT (+/+) and *HB^del/del^* (d/d) embryos transfected with each construct based on the data shown in A. Data represent the mean values ± SE of results obtained from at least three individual explants; n = 9, 3, 7, 4, 7, 6, 4, and 4 for GFP:+/+, JMa-GFP:+/+, JMb-GFP:+/+, ICD-GFP:+/+, GFP:d/d, JMa-GFP:d/d, JMb-GFP:d/d, and ICD-GFP:d/d, respectively. \*\**p* < 0.01; \*\*\**p* < 0.001; \*\*\*\**p* < 0.0001. See also Fig. S5.

## Discussion

In the present study, we propose a model in which dual ErbB signalling with opposing actions is involved in the regulation of mesenchymal cell proliferation. In this model: 1) activation of the heterodimer of ErbB1 and ErbB4 by HB-EGF suppresses cell proliferation, whereas 2) activation of the homodimer of ErbB1 by a certain ErbB1 ligand(s) promotes cell proliferation (Fig. 7). The EGF-ErbB system appears to be subject to complex regulation, producing a range of cellular behaviours in vivo; however, the combinations of EGF ligands and ErbB receptors, and the patterns of ErbB dimerization, that achieve different physiological outcomes have been largely unknown. Only a limited number of studies have investigated these issues. For example, previous studies have demonstrated that HB-EGF contributes to mouse heart muscle homeostasis (Iwamoto et al., 2003) and *trans-retinoic* acid-induced skin hyperplasia (Kimura et al., 2005) through ErbB2/ErbB4 and ErbB1/ErbB2 heterodimers, respectively. HB-EGF and TGFα have been shown to synergistically contribute to skin epithelial sheet migration in mouse eyelid closure (Mine et al., 2005) and alveolar development (Minami et al., 2008) through ErbB1, although the partner of ErbB1 was not identified. NRG1 is involved in heart trabeculation and Schwann cell development through ErbB2/ErbB4 and ErbB2/ErbB3 heterodimers, respectively (Tidcombe et al., 2003). Here, we show for the first time that two ErbB receptor dimer combinations regulate cell proliferation in the same tissue during valvulogenesis in a competitive manner. This provides new insight into the role of ErbB signalling in elaborate regulation of tissue development in vivo.

**Fig. 7.**
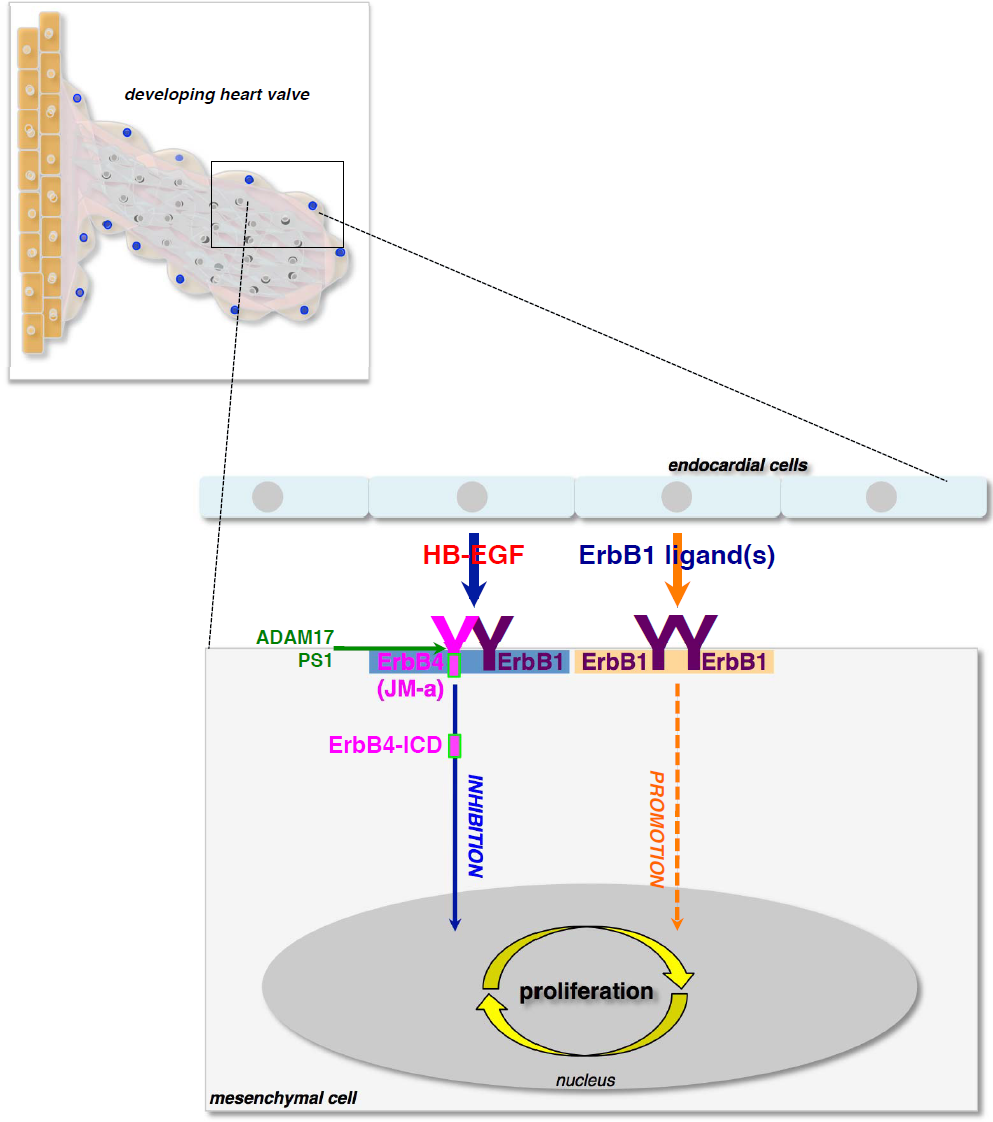
Schematic illustration of the model proposed by this study. A dual ErbB signalling cascade competitively regulates mesenchymal cell proliferation in mouse valvulogenesis as follows: - HB-EGF activates the heterodimer receptor of ErbB1 and ErbB4, resulting in suppression of mesenchymal cell proliferation - Activation of ErbB4 results in its sequential cleavages by ADAM17 and PS1, and ErbB4-ICD nuclear signalling may be involved in this process - An ErbB1 ligand(s) activates the ErbB1 homodimer, resulting in promotion of mesenchymal cell proliferation.

Our results indicate that HB-EGF binds to and activates the heterodimer of ErbB1/ErbB4 in mouse valvulogenesis, and that a certain ErbB1-ligand(s) binds to and activates the homodimer of ErbB1. HB-EGF binds to both ErbB1 and ErbB4. However, why HB-EGF selectively activates the ErbB1/ErbB4 heterodimer and does not activate the ErbB1/ErbB1 homodimer in valvulogenesis is unclear. A key factor underlying this selectivity may be the interaction of HB-EGF with heparan sulfate proteoglycans (HSPGs) through the heparin-binding domain of HB-EGF. The interaction between HB-EGF and HSPGs is essential for normal valvulogenesis, as shown in a mutant knock-in mouse expressing a truncated form of HB-EGF that lacks the heparin-binding domain (*HB^_Δ_*hb*/_Δ_hb^*’) (Iwamoto et al., 2010). It has also been suggested that, in blastocyst implantation, HB-EGF interacts with ErbB4 via HSPGs on the blastocyst cell surface (Paria et al., 1999). Moreover, HSPG-type CD44 is reported to recruit HB-EGF and ErbB4 during remodelling of female reproductive organs (Yu et al., 2002). Based on these findings, we presume that the interaction with HSPGs on the mesenchymal cell surface recruits HB-EGF to the ErbB1/ErbB4 heterodimer specifically, but not to the ErbB1/ErbB1 homodimer.

We found that the ErbB1 homodimer contributes to mesenchymal cell proliferation in developing valves. However, the identity of the ErbB1 ligand(s) that activates the ErbB1 homodimer in the mesenchymal cells, leading to the promotion of cell proliferation, is unknown. Based on our proposed requirement for an interaction between HB-EGF and HSPGs for growth inhibition by the ErbB1/ErbB4 heterodimer, among all ErbB1 ligands (HB-EGF, EGF, TGF-α, ARG, ERG, BTC, and EPG), the ligands with a heparin-binding activity, such as ARG and HB-EGF, may be excluded as candidates for the ErbB1-ligand(s). Moreover, expression of EGF, ARG, ERG, and HB-EGF was detectable by RT-PCR in the cultured cells of WT explants (R. Iwamoto, unpublished observation). Thus, taken together, these data suggest that EGF and ERG may be potential ligands of the ErbB1 homodimer. In addition to HB-EGF, single KO mouse lines of each ErbB1 ligand, including EGF (Luetteke et al., 1999), TGF-α (Luetteke et al., 1999), ARG (Luetteke et al., 1999), ERG (Shirasawa et al., 2004), BTC (Jackson et al., 2003), or EPG (Dahlhoff et al., 2013), have already been established and reported. Furthermore, a double KO mouse line of HB-EGF/BTC (Jackson et al., 2003) and a triple KO mouse line of EGF/TGF-α/ARG (Luetteke et al., 1999) have also been established and reported. However, any abnormalities of valve development in these KO embryos have not been reported so far. These studies strongly suggest that ErbB1 ligands redundantly compensate for each other in terms of their physiological functions in vivo. Thus, to clarify the role of these ligands in this process, it may be necessary to prepare KO mice lacking multiple ErbB1 ligands to show hypoplasia in valve development.

Several lines of evidence have implied that the bone morphogenetic protein (BMP) signalling pathway is also involved in mesenchymal cell proliferation during valvulogenesis. Increases in the activation of Smad 1/5/8 have been reported in hyperproliferating cells of HB-EGF KO valves (Jackson et al., 2003). Moreover, activation of Smad 1/5/8 increases in enlarged valves of TGF-β2 KO mice (Azhar et al., 2009) and CXCR7 KO mice (Sierro et al., 2007; Yu et al., 2011). To understand the entire molecular mechanism governing regulation of cell proliferation in valvulogenesis, it will be important to study the molecular relationship between the ErbB and BMP signalling pathways.

We found that the cleavable JM-a type ErbB4 isoform functions as a receptor for HB-EGF in the suppression of mesenchymal cell proliferation, leading to the possibility of direct nuclear signalling mediated by the ICD of ErbB4 (Fig. 7). In this study, forced expression of cleavable JM-a or ICD rescued ErbB4^−/−^ cells from hyperproliferation, and expression of uncleavable JM-b up-regulated WT cell proliferation by inhibiting HB-EGF-mediated suppression of cell proliferation. Therefore, these results indicate that ErbB4 cleavage is essential, and that the ErbB4 ICD is necessary and sufficient for the inhibitory signalling in valve mesenchymal cells. Several lines of in vitro evidence indicate that the biological functions of JM-a and JM-b are different (Carpenter, 2003; Veikkolainen et al., 2011). However, the physiological functions of each isoform in vivo are unclear. Thus far, only one report has directly indicated a physiological function of JM-a, demonstrating a role in astrocyte differentiation during neural development (Sardi et al., 2006). Thus, our study adds valvulogenesis as a novel physiological process in which JM-a physiologically functions in vivo. The ErbB4-ICD form, which is an artificially truncated transmembrane domain, has been thought to be constitutively active (Linggi et al., 2006; Muraoka-Cook et al., 2009). However, the rescue activity of the ICD in *ERBB4^−/−^* explants was weaker than that of the full-length JM-a form. A similar result was obtained in our experiment with *HB^del/del^* explants. These results suggest that, for

ErbB4-ICD to fully function in nuclear signalling, the ICD must be synthesised, presented on the cell surface as the full-length transmembrane JM-a type of ErbB4, and then sequentially cleaved on the cell surface and released into the cytoplasm.

ADAM17 (also known as tumour necrosis factor-α converting enzyme; TACE) is a member of the ADAM family metalloproteases (Scheller et al., 2011) and is suggested to be a major sheddase of HB-EGF during valvulogenesis because *ADAM17* KO valves phenocopy *HBEGF* KO valves (Jackson et al., 2003; Wilson et al., 2013). Similar to HB-EGF, ectodomain shedding of JM-a type ErbB4 is also thought to be mediated by ADAM17 (Rio et al., 2000; Veikkolainen et al., 2011). Shedding of ErbB4 by ADAM17 is thought to be required before proceeding with intramembrane proteolysis by PS1, which results in releasing the ErbB4 ICD into the cytoplasm (Ni et al., 2001; Veikkolainen et al., 2011). Therefore, in the inhibitory process mediated by the HB-EGF-ErbB4 axis, ADAM17 may act as a sheddase with dual functions to shed HB-EGF in endocardial cells and ErbB4 in mesenchymal cells. The molecular mechanism governing the regulation of this protease during valve remodelling processes should be investigated further in future studies.

HB-EGF plays important roles in cancer cell proliferation, malignancy, metastatic potential, and chemotherapy resistance (Miyamoto et al., 2006). In particular, HB-EGF expression is increased in advanced ovarian cancer compared with normal ovarian tissue (Miyamoto et al., 2004) and associated with poor clinical outcomes (Tanaka et al., 2005). HB-EGF is not only expressed in cancer cells, but also in the cancer-surrounding stroma, which is involved in tumour progression (Murata et al., 2011). Thus, HB-EGF is recognised as a possible target for cancer therapy. An anti-HB-EGF antibody (Miyamoto et al., 2011) and CRM197 (a nontoxic mutant form of diphtheria toxin that neutralises HB-EGF activity) are undergoing clinical development as anticancer drugs (Nam et al., 2014). However, such blocking therapies also have potential side effects associated with the inhibition of “physiological” HB-EGF. Interestingly, such physiological processes that promote cell proliferation have not been found so far (Mekada and Iwamoto, 2008; Iwamoto and Mekada, 2012). Conversely, in valvulogenesis, HB-EGF functions as an “inhibitory” factor. This suggests the concept of inhibiting the proliferation of cancer cells expressing HB-EGF by converting the HB-EGF function in proliferation from “promotion” to “suppression”. This could be achieved by introducing a key factor or system into the cancer cells, which governs the inhibitory mechanism in mesenchymal cells during valvulogenesis. The present study provides us with hope that ErbB4 or the pathway downstream of ErbB4 will be a strong candidate for such a key factor. Further study of the regulatory mechanism mediated by the HB-EGF-ErbB4 signalling axis in valvulogenesis may shed light on establishing a novel method for molecular targeted cancer therapy.

## Materials and methods

### Mice

The generation of *HBEGF-KO* (*HB^del/del^*) mice has been described previously (Iwamoto et al., 2003). These mice were backcrossed for more than 13 generations onto a C57BL/6J background. *Waved-2* mice (Luetteke et al., 1994) were purchased from the Jackson Laboratory. *Waved-2* mice with a mixed background of C57BL/6JEi and C3H/HeSnJ when purchased were backcrossed for more than 5 generations onto a C57BL/6J background. To establish double mutant mice of *waved-2* and *HB^del^, HB^del/+^* mice were mated with *waved-2* heterozygous (*wa2/+*) mice. *ERBB4-KO* (Gassmann et al., 1995) mice rescued from embryonic lethality by cardiac expression of HER4 (*ERBB4^−/−^ HER4^heart^*) (Tidcombe et al., 2003) were kindly donated by E. Clark (MRC National Institute for Medical Research) and used under the kind permission of M. Gassmann (University of Basel). *ERBB4^+/−^ HER4^heart^* males were crossed with *ERBB4^+/−^ HER4^heart^* females to generate *ERBB4^−/−^* (null)-, *ERBB4^+/−^ HER4^heart^*, and WT embryos. The discovery of a vaginal plug was considered as E0.5. To analyse valve development and ex vivo cultures, timed pregnant females were sacrificed, and embryos at a suitable stage for each analysis as shown below were dissected from the uteri and placed in phosphate-buffered saline (PBS). All mice used in this study were housed in the Animal Care Facility of Osaka University Research Institute for Microbial Diseases according to the Institutional guidelines for laboratory animals. The use and treatment of animals was approved by the Institutional Biosafety Committee on Biological Experimentation and the Institutional Animal Experimentation Committee at Osaka University.

### Plasmids

Full-length cDNA encoding human ErbB1 was kindly provided by M. Shibuya (The University of Tokyo). Full-length cDNA encoding human ErbB2 was kindly provided by K. Maruyama (The University of Tokyo). Full-length cDNAs encoding human ErbB3 and JM-b CYT-1 type ErbB4 were kindly provided by K. Elenius (University of Turku). Myc-tagged dn-ErbB1 (Kashles et al., 1991), myc-tagged dn-ErbB2 (Jones and Stern, 1999), myc-tagged dn-ErbB3 (Ram et al., 2000), and myc-tagged dn-ErbB4 (Jones et al., 1999) were generated by PCR using cDNA encoding each corresponding full-length cDNA as a template. Myc-tagged JM-a CYT-1 type ErbB4 and myc-tagged CYT-1 type ErbB4-ICD (Sundvall et al., 2007) were generated by site-directed mutagenesis via PCR using cDNA encoding full length JM-b CYT-1 type ErbB4 as a template. All generated constructs were cloned into pLVSIN-IRES ZsGreen1 (Clontech) to produce lentiviruses for infection of cushion explant cultures as described below. Preparation and titering of lentiviruses were carried out according to the manufacturer’s instructions. ZsGreen1 is an analogue of green fluorescent protein (GFP) and referred to as “GFP” in this study.

### Endocardial cushion explant cultures

To examine the proliferation of valve mesenchymal cells in vitro, the original method of endocardial cushion explant culture (Camenisch et al., 2002) was modified as described previously (Iwamoto et al., 2010). In brief, a cushion explant from an E10.5 embryonic heart was placed on a type I collagen gel (Cellmatrix Type I-A, Nitta Gelatin Co.) containing 0.5 mg/ml hyaluronic acid (HA-Col; 0.5 ml/well in a 24-well plate), with one side of the endocardium in contact with the gel surface. The explant was grown in 0.5 ml Medium 199 supplemented with 1% fetal calf serum (FCS), 100 U/ml penicillin, 100 μg/ml streptomycin, and 1% Insulin-Transferrin-Selenium-X (GIBCO BRL) at 37°C in 5% CO_2_. During the culture period, endocardial cells from the endocardium of the cushion explants (ON-gel cells) proliferate and spread on the surface of the HA-Col, while differentiated mesenchymal cells from the endocardium of the explants (IN-gel cells) invade into the HA-Col and proliferate. In experiments using the EGFR inhibitor ZD1839 (AstraZeneca) or recombinant human soluble HB-EGF protein (R&D), these were added on day 5 after starting the culture, and then the cells were cultured for a further 5 days. After a total of 10 days in culture, the proliferation of ON-gel and IN-gel cells was measured by BrdU incorporation. An image of the anti-BrdU polyclonal antibody (Abcam)-stained explant was captured by a confocal laser scanning microscope system (LSM5 PASCAL, Carl Zeiss) and analysed with Adobe Photoshop software.

In experiments using lentivirus expression vectors encoding the cDNAs described above, lentivirus infection was carried out on day 5 after starting the culture, and then the cells were cultured for a further 5 days. To infect both IN-gel and ON-gel cells, 10–20 μl of concentrated virus solution (>1 × 10^9^ TU/ml) in Medium 199 was directly infused into the gel beneath the explant via an ultrathin capillary. After a total 10 days in culture, the proliferation of ON-gel and IN-gel cells was measured by immunocytochemistry with a polyclonal antibody against Ki67 (Novocastra) as a mitotic marker. An image of the Ki67-stained explant was captured by the confocal laser scanning microscope system and analysed with Adobe Photoshop software. In these cDNA introduction experiments, the total colour of the captured image was automatically converted to clearly distinguish signals of GFP and Ki67 (Fig. 3B). All data were analysed in a blinded manner.

### Histological analysis

Haematoxylin and eosin staining and measurement of valve size were performed as described previously (Iwamoto et al., 2003). At E18.5 to postnatal day 0, the hearts of embryos or pups were fixed with 4% paraformaldehyde and embedded in optimal cutting temperature compound. For immunohistochemistry to detect phosphorylated ErbB1, serial frozen sections (8 μm) were incubated with an anti-pY1068-ErbB1 rabbit monoclonal antibody (Cell Signaling) using the Duolink II system (Sigma-Aldrich) according to the manufacturer’s instructions. For immunohistochemistry to detect total ErbB1, sections were incubated with a biotinylated anti-ErbB1 rabbit polyclonal antibody (Cell Signaling) and then Alexa488-conjugated streptavidin (Molecular Probes).

### RT-PCR analysis of ErbB receptors

Total RNA from cushion explant cells (mixture of IN-gel and ON-gel cells after removal of explants) cultured for 10 days was isolated using an RNeasy Micro kit (Qiagen). Total RNA was reverse transcribed using reverse transcriptase ReverTra Ace (TOYOBO).

PCR analysis was performed using ExTaq (TOYOBO) for each gene as follows:

Mouse *ERBB1* (94°C for 30 sec, 57°C for 30 sec, 72°C for 60 sec; 40 cycles), mouse *ERBB2* (94°C for 30 sec, 67°C for 30 sec, 72°C for 60 sec; 40 cycles), mouse *ERBB3* (94°C for 30 sec, 67°C for 30 sec, 72°C for 60 sec; 40 cycles), mouse *ERBB4_JM-a&b* type (94°C for 30 sec, 63°C for 30 sec, 72°C for 60 sec; 40 cycles), mouse *ERBB4_JM-a* type specific (94°C for 30 sec, 63°C for 30 sec, 72°C for 60 sec; 40 cycles), and mouse *ERBB4_JM-b* type specific (94°C for 30 sec, 60°C for 30 sec, 72°C for 60 sec; 40 cycles).

The gene-specific primers were as follows:

Mouse *ERBB1* (5′ - TGGAGGAAAAGAAAGTCTGCCAAGGCAC -3′ and 5′ - ACATCCATCTGATAGGTGGTGGGGTT -3′), mouse *ERBB2* (5′ - GGGGGAGCTGGTCGATGCTG -3′ and 5′ - CTTTGGTTACCCCCACTGCC -3′), mouse *ERBB3* (5′ - GGGTCGCGAACAGTTCTCCC -3′ and 5′ - GAGCGGGGTGACGGGAGTAA -3′), mouse *ERBB4_JM-a&b* type (5′ - GAAATGTCCAGATGGCCTACAGGG -3′ and 5′ - CTTTTTGATGCTCTTTCTTCTGAC -3′), mouse *ERBB4_JM-a* type specific (5′ - AACGGTCCCACTAGTCATGACTGC -3′ and 5′ - TGGATAGTGGGACTCAGACACACT -3′), and mouse *ERBB4_JM-b* type specific (5′ - ATAGGTTCAAGCATTGAAGACTGC -3′ and 5′ - TGGATAGTGGGACTCAGACACACT -3′).

### Immunoblotting

Human HEK293 cells were transiently transfected with vectors encoding JM-a, JM-b, or ICD by Lipofectamine 2000 (Invitrogen) according to the manufacturer’s instructions, and cultured for 36 h in Dulbecco’s modified Eagle’s medium (DMEM) containing 10% heat-inactivated FCS. Then, the cells were washed and the medium was replaced with fresh medium with or without 10% FCS. After overnight culture, the cells were treated in the presence or absence of 100 nM TPA for 30 min, lysed with cell lysis buffer [60 mM 1-O-n-octyl-β-D-glucopyranoside, 0.15 M NaCl, 10 mM HEPES-NaOH, pH 7.2, 1/100× protease inhibitor cocktail (Nacalai)], and centrifuged at 20,000 × *g* for 30 min. The supernatant was collected, mixed with sodium dodecyl sulfate (SDS) gel sample buffer with or without 10% β-mercaptoethanol, and loaded onto 7.5% SDS electrophoresis gels. Separated proteins were electrotransferred onto a polyvinylidene difluoride membrane (Immobilon-P, Millipore). The membrane was blocked in TTBS (10 mM Tris-HCl, pH 7.5, 0.15 M NaCl, 0.05% Tween 20) containing 1% dry skim milk for 1 h, followed by incubation with a mouse anti-myc (9E10) monoclonal antibody (Santa Cruz Biotechnology) at 1 μg/ml in TTBS for 1 h. The membrane was then incubated with a horseradish peroxidase-conjugated secondary antibody (1:3000) for 1 h. Chemiluminescent detection was carried out with the ECL detection system (Pierce) according to the manufacturer’s instructions.

### Immunostaining

Human HT1080 cells were transiently transfected with lentiviruses containing each construct and then cultured for 72 h in DMEM containing 10% heat-inactivated FCS. Then, the cells were washed and the medium was replaced with fresh medium without FCS. After overnight culture, the cells were fixed with 4% paraformaldehyde in PBS at room temperature for 15 min, permeabilized with 0.5% Triton X-100 in PBS at room temperature for 15 min, blocked with blocking solution (2% donkey serum, 1% bovine serum albumin, and 0.05% Tween 20 in PBS), and incubated with a mouse anti-myc (9E10) monoclonal antibody (Santa Cruz Biotechnology) at 2 μg/ml in blocking solution at 4°C overnight. The cells were then incubated with Alexa546-conjugated donkey anti-mouse IgG (1:500; Molecular Probes).

### Data analysis

All quantitative data are presented as the mean ± SE. Statistical significance was assessed by the two-tailed Student’s t-test. A value of *p* < 0.05 was considered statistically significant.

## Acknowledgements

We thank M. Hamaoka, K. Nishiwaki, Y. Numata, and T. Yoneda for technical assistance.

## Author Contributions

R.I. designed the study, performed most experiments, analysed all data, and wrote the manuscript. N.M. performed the histology of *HB^del^* and *waved-2* double mutant valves, and analysed data of the size measurements. H.M. constructed plasmids encoding several ErbB4 variants. E.M. supervised the study and wrote the manuscript. All authors discussed the results and commented on the manuscript.

## Supplementary Information

Supplementary Information includes five Supplemental Figures is available.

